# ABlooper: Fast accurate antibody CDR loop structure prediction with accuracy estimation

**DOI:** 10.1101/2021.07.26.453747

**Authors:** Brennan Abanades, Guy Georges, Alexander Bujotzek, Charlotte M. Deane

## Abstract

Antibodies are a key component of the immune system and have been extensively used as biotherapeutics. Accurate knowledge of their structure is central to understanding their antigen binding function. The key area for antigen binding and the main area of structural variation in antibodies is concentrated in the six complementarity determining regions (CDRs), with the most important for binding and most variable being the CDR-H3 loop. The sequence and structural variability of CDR-H3 make it particularly challenging to model. Recently deep learning methods have offered a step change in our ability to predict protein structures. In this work we present ABlooper, an end-to-end equivariant deep-learning based CDR loop structure prediction tool. ABlooper rapidly predicts the structure of CDR loops with high accuracy and provides a confidence estimate for each of its predictions. On the models of the Rosetta Antibody Benchmark, ABlooper makes predictions with an average CDR-H3 RMSD of 2.49Å, which drops to 2.05Å when considering only its 76% most confident predictions.

## 1 Introduction

### 1.1 Antibody Structure

Antibodies are a class of protein produced by B-cells during an immune response. Their ability to bind with high affinity and specificity to almost any antigen, makes them attractive for use as therapeutics [1].

Knowledge of the structure of antibodies is becoming increasingly important in biotherapeutic development [2]. However, experimental structure determination is time-consuming and expensive so it is not always practical or even possible to use routinely. Computational modelling tools have allowed researchers to bridge this gap by predicting large numbers of antibody structures to a high level of accuracy [3, 4]. For example, models of antibody structures have recently been used for virtual screening [5] and to identify coronavirus-binding antibodies that bind the same epitope with very different sequences [6].

The overall structure of all antibodies is similar and therefore can be accurately predicted using current methods (e.g. [3]). The area of antibodies that it is hardest to model are the sequence variable regions that provide the structural diversity necessary to bind a wide range of antigens. This diversity is largely focused in six loops known as the complementarity determining regions (CDRs). The most diverse of these CDRs and therefore the hardest to model is the third CDR loop of the heavy chain (CDR-H3) [7].

### 1.2 Deep Learning for Protein Structure Prediction

At CASP14 [8], DeepMind showcased AlphaFold2 [9], a neural network capable of accurately predicting many protein structures. The method relies on the use of equivariant neural networks and an attention mechanism. More recently, RoseTTAFold, a novel neural network based on equivariance and attention was shown to obtain results comparable to those of AlphaFold2 [10].

These methods both rely on the use of equivariant networks. For a network to be equivariant with respect to a group, it must be able to commute with the group action. For rotations, this means that rotating the input before feeding it into the network will have the same result as rotating the output. In the case of proteins, using a network equivariant to both translations and rotations in 3D space allows us to learn directly from atom coordinates. This is in contrast to previous methods like TrRosetta [11] or the original version of AlphaFold [12] that predict invariant features, such as inter-residue distances and orientations which are then used to reconstruct the protein. A number of approaches for developing equivariant networks have been recently developed (e.g. [13]).

In this paper we explore the use of an equivariant approach to CDR structure prediction. We chose to use E(n)-Equivariant Graph Neural Networks (E(n)- EGNNs) [14] as our equivariant approach due to their speed and simplicity.

### 1.3 Deep Learning for Antibody Structure Prediction

Deep learning based approaches have also been shown to improve structure prediction in antibodies, for example DeepAb [4], was shown to outperform all currently available antibody structure prediction methods. DeepAb is similar to TrRosetta and the original version of AlphaFold in that deep learning is used to obtain inter-residue geometries that are then fed into a computationally expensive energy minimisation method to produce the final structure.

In this work we present ABlooper, a fast and accurate tool for antibody CDR loop structure prediction. By leveraging E(n)-EGNNs, ABlooper directly predicts the structure of CDR loops. By simultaneously predicting multiple structures for each loop and comparing them amongst themselves, ABlooper is capable of estimating a confidence measure for each predicted loop.

## 2 Methods

### 2.1 Data

The data used to train, test and validate ABlooper was extracted from SAbDab [15], a database of all antibody structures contained in the PDB [16]. Structures with a resolution better than 3.0Å and no missing backbone atoms within any of the CDRs were selected. The CDRs were defined using the IMGT numbering scheme [17].

For easy comparison with different pipelines, we used the 49 antibodies from the Rosetta Antibody Benchmark as our test set. For validation, 100 structures were selected at random. It was ensured that there were no structures with the same CDR sequences in the training, testing and validation sets. Sequence redundancy was allowed within the training set to expose the network to the existence of antibodies with identical sequence but different structural conformations. This resulted in a total of 3438 training structures.

Additionally, we use a secondary test set composed of 114 antibodies (SAbDab Latest Structures) with a resolution of under 2.3Å and a maximum CDR-H3 loop length of 20, which were added to SAbDab after the initial test, train and validation sets were extracted (8^*th*^November 2020 − 24^*th*^May 2021). A list containing the PDB IDs of all the structures used in the train, test, and validation sets is given in the SI.

ABodyBuilder was used to build models of all the structures. Structural models were generated using the singularity version of ABodyBuilder [3] (fragment database from 8^*th*^July 2021) excluding all templates with a 99% or higher sequence identity. ABlooper CDR models for the test sets were obtained by remodelling the CDR loops on ABodyBuilder models.

### 2.2 Deep Learning

ABlooper is composed of five E(n)-EGNNs, each one with four layers, all trained in parallel. The model is trained on the position of the C_*α*_-N-C-C_*β*_ backbone atoms for all six CDR loops plus two anchor residues at either end. An input geometry is generated by flattening each of the CDR loops while leaving the anchor residues in place. The model is given four different types of features per residue resulting in a 41-dimensional vector. These include a one-hot encoded vector describing the amino acid type, the atom type and which loop the residue belongs to. Additionally, sinusoidal positional embeddings are given to each residue describing how close it is to the anchors.

Each of the five networks is trained on antibody crystal structures to independently predict the loop conformation for all of the CDRs. The RMSD between the prediction and true structure is used as the loss function. The output from the five networks is then averaged to obtain a final prediction. To make the final prediction physically plausible, an L1 loss between all intraresidue atom distances is used. More details on the implementation and training of ABlooper can be found in the SI.

### 2.3 Loop relaxation

During training, ABlooper is encouraged to predict physically plausible CDR loops via the intra-residue atom distance loss term. However, ABlooper occasionally produces loops with incorrect backbone geometries. To enforce correct backbone geometries we relax the loop using the Amber14 [18] protein force field. This relaxation step typically results in a small loss in accuracy, but ensures that predicted loops are physically plausible. More details on how this is implemented can be found in the SI.

### 2.4 DeepAb and AlphaFold2

Structural models for DeepAb were generated using the open source version of the code (available at https://github.com/RosettaCommons/DeepAb). As suggested in their paper [4], we generated five decoys per structure. This took around 10 minutes per antibody on an 8-core Intel i7-10700 CPU.

Antibody structures were generated using the open source version of AlphaFold2 (available at https://github.com/deepmind/alphafold). We used the “full dbs” preset and allowed it to use templates from before the 14^*th*^May 2020. As AlphaFold2 is intended to predict single chains [9], we predicted and aligned the heavy and light chain independently before comparing to other methods. On a 20-core Intel 6230 CPU this took around 3 hours per antibody modelled.

## 3 Results

### 3.1 Using ABlooper to predict CDR loops on modelled antibody structures

We used ABlooper to predict the CDRs on ABodyBuilder models of the Rosetta Antibody Benchmark (RAB) and the SAbDab Latest Structures (SLS) sets. The RMSD between the C_*α*_-N-C-C_*β*_ atoms in the backbone of the crystal structure and the predicted CDRs for both test sets is shown in Table 1.

**Table 1:** Performance comparison between AlphaFold2, ABodyBuilder, DeepAb and ABlooper for both test sets.The mean RMSD to the crystal structure across each test set for the six CDRs is shown. RMSDs for each CDR are calculated after superimposing their corresponding chain to the crystal structure. RMSDs are given in Angstroms (Å). (*) It is likely that AlphaFold2 used at least some of the structures in the benchmark set during training. Similarly, structures in the SAbDab Latest Structures set may have been used for training DeepAb.

| Method | CDR-H1 | CDR-H2 | CDR-H3 | CDR-L1 | CDR-L2 | CDR-L3 |
| --- | --- | --- | --- | --- | --- | --- |
| Rosetta Antibody Benchmark |  |  |  |  |  |  |
| AlphaFold2* | 0.84 | 0.99 | 2.87 | 0.53 | 0.49 | 0.95 |
| ABodyBuilder | 1.08 | 0.99 | 2.77 | 0.69 | 0.50 | 1.12 |
| DeepAb | 0.83 | 0.93 | 2.44 | 0.50 | 0.44 | 0.85 |
| ABlooper | 0.92 | 1.01 | 2.49 | 0.62 | 0.52 | 0.97 |
| ABlooper Unrelaxed | 0.90 | 1.03 | 2.45 | 0.61 | 0.51 | 0.93 |
| SAbDab Latest Structures |  |  |  |  |  |  |
| ABodyBuilder | 1.24 | 1.07 | 3.25 | 0.88 | 0.57 | 1.03 |
| DeepAb* | 1.00 | 0.82 | 2.49 | 0.59 | 0.45 | 0.90 |
| ABlooper | 1.14 | 0.97 | 2.72 | 0.74 | 0.55 | 1.04 |
| ABlooper Unrelaxed | 1.14 | 0.99 | 2.66 | 0.73 | 0.54 | 1.01 |

ABlooper achieves lower mean RMSDs than AbodyBuilder for most CDRs (Table 1). By far the largest improvement is for the CDR-H3 loop, where due to the large structural diversity, homology modelling performs worst [3]. ABlooper predicts loops of a similar accuracy to AlphaFold2 and DeepAb for all CDRs except CDR-H3, where ABlooper and DeepAb outperform AlphaFold2.

One potential source of error for ABlooper are the model frameworks generated by ABodyBuilder, so we examined its resilience to the small deviations seen in these models and found little to no correlation between framework error and CDR prediction error (see SI).

### 3.2 Prediction diversity as a measure of prediction quality

ABlooper predicts five structures for each loop. We found that the average RMSD between predictions can be used as a measure of certainty of the final averaged prediction. If all five models agree on the same conformation, then it is more likely that it will be the correct conformation, if they do not, then the final prediction is likely to be less accurate (Figure 1). This allows ABlooper to give a confidence score for each predicted loop. As shown in Figure 1D, this score can be used as a filter, removing structures which are expected to be incorrectly modelled by ABlooper. For example, by setting a 1.5Å inter-prediction RMSD cut-off on structures from the Rosetta Antibody Benchmark, the average CDR-H3 RMSD for the set can be reduced from 2.49Å to 2.05Å while keeping 76% of predictions. As expected, accuracy filtering has a tendency to remove longer CDR-H3 predictions but it is not exclusively correlated to length (see SI).

**Figure 1:**
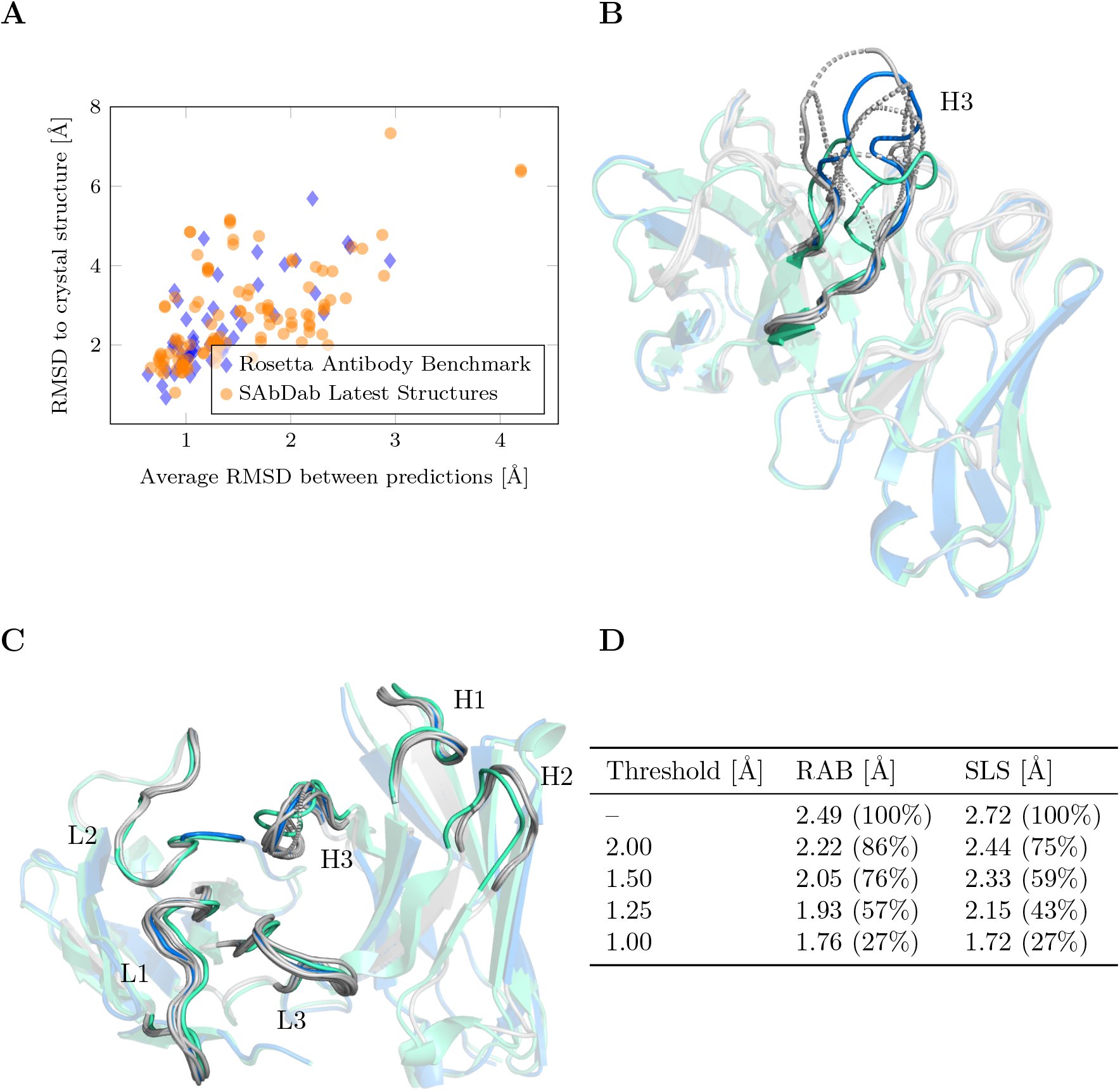
(**A**) CDR-H3 loop RMSD between final averaged prediction and crystal structure compared to average RMSD between the five ABlooper predictions for both the Rosetta Antibody Benchmark and the SAbDab Latest Structures set. (**B**) An example of a poorly predicted CDR-H3 loop. All five predictions are given in grey, with the final averaged prediction in blue and the crystal structure in green. The predictions from the five networks are very different, indicating an incorrect final prediction. (**C**) Example of correctly predicted CDR loops. All five predictions are similar, indicating a high confidence prediction. (Colours are the same as in (B)) (**D**) Effect of removing structures with a high CDR-H3 inter-prediction RMSD on the averaged RMSD for the relaxed set. The number of structures remaining after each quality cut-off is shown as a percentage. Data shown for the Rosetta Antibody Benchmark (RAB) and the SAbDab Latest Structures (SLS) sets.

## 4 Discussion

We present ABlooper, a fast and accurate tool for predicting the structures of the CDR loops in antibodies. It builds on recent advances in equivariant graph neural networks to improve CDR loop structure prediction. By predicting each loop multiple times, it is also capable of producing an accuracy estimate for each generated loop structure.

On an NVIDIA Tesla V100 GPU, the unrelaxed version of ABlooper can predict the CDR backbone atoms for one hundred structures in under five seconds. Loop relaxation and side-chain prediction are the most computationally expensive parts of the pipeline taking around ten seconds per structure. ABlooper outperforms ABodyBuilder (a state of the art homology method) and produces antibody models of similar accuracy to both AlphaFold2 and DeepAb, but on a far faster timescale. The model used for ABlooper is available at: https://github.com/oxpig/ABlooper.

## Supporting information

Train and test datasets

Train and test datasets

## 5 Funding

This work was funded by the Engineering and Physical Sciences Research Council (EPSRC) with grant number: EP/S024093/1.

## Notes

https://github.com/oxpig/ABlooper

### Competing Interest Statement

The authors have declared no competing interest.

### Summary of Updates

Added a loop relaxation step to correct for abnormal geometries.

