## Supplementary Information for "ABlooper: Fast accurate antibody CDR loop structure prediction with accuracy estimation"

#### A Network implementation and training details

ABlooper was implemented in Pytorch [1]. The model is composed of five E(n)-EGNNs [2] all simultaneously predicting the structure of each loop. The implementation of E(n)-EGNN is based on the open source version of Phil Wang [3]. The main difference between our implementation of E(n)-EGNN and the original is the normalisation of relative coordinates before each coordinate update. This was found to help stability during training. Additionally, edge features were not used in our model. With these changes, the equations describing the E(n)-EGNN algorithm become:

$$\begin{aligned}
\mathbf{r_{ij}} &= \mathbf{x_i^l} - \mathbf{x_j^l} \\
\mathbf{m_{ij}} &= \phi_e(\mathbf{h_i^l}, \mathbf{h_j^l}, \|\mathbf{r_{ij}}\|^2) \\
\mathbf{x_i^{l+1}} &= \mathbf{x_i^l} + C \sum_{j \neq i} \hat{\mathbf{r}}_{\mathbf{ij}} \, \phi_x(\mathbf{m_{ij}}) \\
\mathbf{m_i} &= \sum_j \mathbf{m_{ij}} \\
\mathbf{h_i^{l+1}} &= \mathbf{h_i^l} + \phi_h(\mathbf{h_i^l}, \mathbf{m_i})
\end{aligned}$$

Where $\mathbf{x_i^l}$ are the coordinates of node $i$ after layer $l$ and $\mathbf{h_i^l}$ is the 41-dimensional feature vector of node $i$. In this notation, $\phi$ is a Multi Layer Perceptron (MLP), $C$ represents a learnable parameter at each layer and $\hat{\mathbf{r}}_{\mathbf{ij}}$ is the normalised distance between the points $i$ and $j$.

The edge MLP ($\phi_e$), is composed of Linear-SiLU-Linear-SiLU layers, yielding 32 features for each pair of nodes. The node and coordinate update MLPs ($\phi_x$ and $\phi_h$) are both composed of Linear-SiLU-Linear layers.

Two different losses were used during training. To quantify the structural similarity between the predicted and true structures, RMSD was used. To ensure distances between neighbouring atoms in the chain were conserved, an L1-loss between the true and predicted inter-atom distances was used. This was composed of five terms between the following pairs of atoms: $C_\alpha^i$-$C_\alpha^{i+1}$, $C_\alpha^i$-$C_\beta^i$, $C_\alpha^i$-$N^i$, $C_\alpha^i$-$C^i$, $C^i$-$N^{i+1}$.

Each of the five E(n)-EGNNs were trained to make predictions independently by minimising the RMSD between their prediction and the crystal structure. To ensure that the final combined prediction of all E(n)-EGNNs was physically plausible, the L1-loss was used on the final averaged structure.

The model was trained in two phases. First it was trained with early stopping without the L1-loss term using the RAdam [4] optimiser with a learning rate of $10^{-3}$ and a weight decay of $10^{-3}$. In the second stage, the L1-loss term was added with a weighting of 1.0. For this stage, the model was trained using the Adam optimiser with a learning rate of $10^{-4}$ and early stopping. In total the model was trained for around 200 epochs for the first stage and 150 for the second. The average RMSD of predicted structures increased around 10% in the second stage of training. However, after the second stage of training all generated structures had physically plausible inter-atomic distances.

### B Loop relaxation and side-chain prediction

ABlooper will occasionally generate loops with physically implausible backbone geometries. To fix this, we model the position of side-chain atoms and relax the predicted loops using a restrained energy minimisation procedure. As our energy function, we use the AMBER14 [5] protein force field with an additional harmonic potential term keeping the positions of backbone atoms close to their original positions. The spring constant of the harmonic potential is set to 10 kcal/mol$Å^2$. Energy minimisation is done using the Langevin Integrator in the OpenMM python package [6].

### C Prediction dependence on framework quality

Here we explore whether the quality of the frameworks generated by ABodyBuilder affect the quality of ABlooper's CDR-H3 loop predictions. SI Figure 1 shows that, ABlooper is capable of consistently predicting accurate CDR-H3 loop structures independent of small errors in ABodyBuilder framework predictions.

### D Prediction diversity against loop length

Longer CDR loops have been shown to be harder to model [7, 8]. As shown in SI Figure 2, longer loops tend to have higher prediction diversity. This means that accuracy filtering will tend to select shorter loops. However, we find that filtering by prediction diversity is effective even amongst loops of the same length. For example, if we select the 75% best scoring CDR-H3 predictions for each loop length separately, the average RMSD over both test sets drops from 2.65Å to 2.45Å.

### E Prediction diversity as a measure of prediction quality for all CDR loops

As shown in Figure 1 of the manuscript, the accuracy of a CDR-H3 loop predicted by ABlooper can be estimated by the diversity of predicted conformations. In SI Figure 3, we show this relation for each of the CDR loops. In contrast to CDR-H3, the other CDR loops have a limited number of canonical

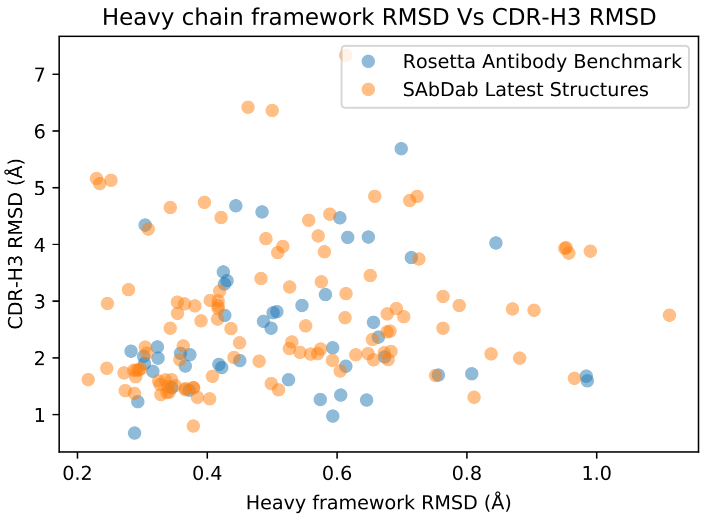


SI Figure 1: Heavy chain framework RMSD of the ABodyBuilder model against CDR-H3 RMSD of ABlooper Predictions. Framework RMSDs were calculated after superimposing the predicted ABodyBuilder framework onto the crystal structure.

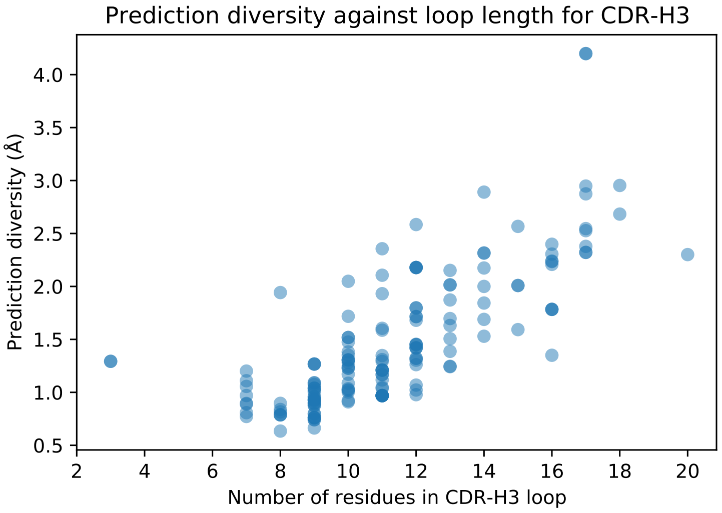


SI Figure 2: Prediction diversity RMSD against the number of amino acids in CDR-H3 loop for all of the structures in the RAB and the SLS datasets.

forms [9] and because of this, the distributions for other CDRs follow a less clear trend than for CDR-H3.

Based on these plots, we explored a few cases in more detail. For example, in the case of the antibody with PDB code 3mlr, the CDR-L3 loops is predicted incorrectly (RMSD $\approx$ 8.6Å) with what could be considered an acceptable level of confidence. When looking at the predicted structure, it was found that this is due to ABodyBuilder incorrectly modelling the framework region. This shows the reliance of ABlooper on obtaining accurate templates for the framework.

A second case is the model of the antibody with PDB code 3lmj, for which ABlooper obtains an RMSD of around 6.3Å for the CDR-H2 loop. In this example, the model incorrectly predicts an interaction between the CDR-H3 and CDR-H2 loops, displacing the CDR-H2. In this case the predicted accuracy is low.

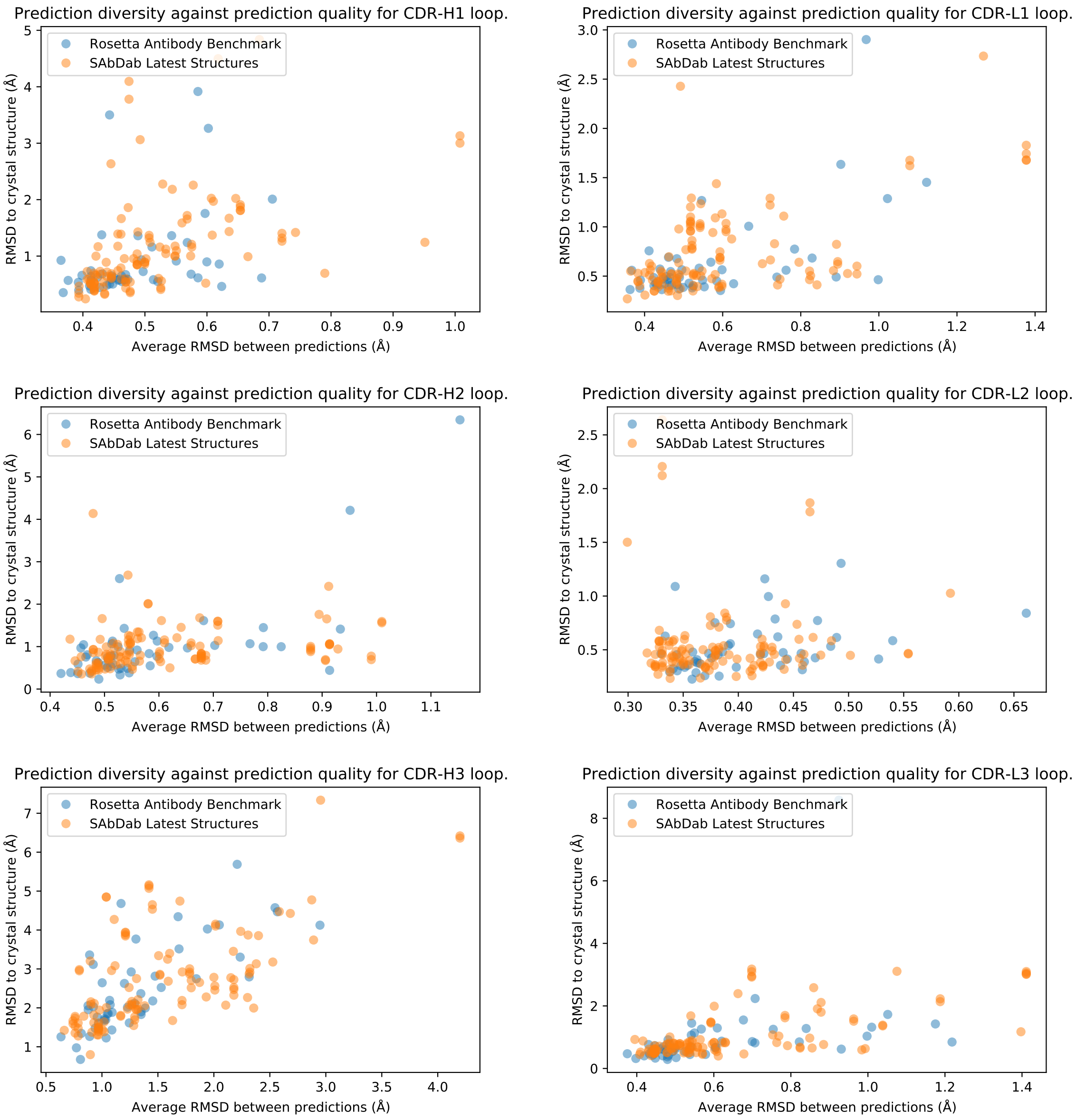


SI Figure 3: Prediction diversity against prediction quality for each of the six CDR loops.

### References

[1] Adam Paszke, Sam Gross, Francisco Massa, Adam Lerer, James Bradbury, Gregory Chanan, Trevor Killeen, Zeming Lin, Natalia Gimelshein,

Luca Antiga, Alban Desmaison, Andreas Kopf, Edward Yang, Zachary DeVito, Martin Raison, Alykhan Tejani, Sasank Chilamkurthy, Benoit Steiner, Lu Fang, Junjie Bai, and Soumith Chintala. Pytorch: An imperative style, high-performance deep learning library. In H. Wallach, H. Larochelle, A. Beygelzimer, F. d'Alché-Buc, E. Fox, and R. Garnett, editors, *Advances in Neural Information Processing Systems 32*, pages 8024–8035. Curran Associates, Inc., 2019.

[2] Victor Garcia Satorras, Emiel Hoogeboom, and Max Welling. E (n) equivariant graph neural networks. *arXiv preprint arXiv:2102.09844*, 2021.

[3] Phil Wang and Eric Alcaide. EGNN - Pytorch. `https://github.com/lucidrains/egnn-pytorch`, 2021.

[4] Liyuan Liu, Haoming Jiang, Pengcheng He, Weizhu Chen, Xiaodong Liu, Jianfeng Gao, and Jiawei Han. On the variance of the adaptive learning rate and beyond. In *Proceedings of the Eighth International Conference on Learning Representations (ICLR 2020)*, April 2020.

[5] James A Maier, Carmenza Martinez, Koushik Kasavajhala, Lauren Wickstrom, Kevin E Hauser, and Carlos Simmerling. ff14sb: improving the accuracy of protein side chain and backbone parameters from ff99sb. *Journal of chemical theory and computation*, 11(8):3696–3713, 2015.

[6] Peter Eastman, Jason Swails, John D Chodera, Robert T McGibbon, Yutong Zhao, Kyle A Beauchamp, Lee-Ping Wang, Andrew C Simmonett, Matthew P Harrigan, Chaya D Stern, et al. Openmm 7: Rapid development of high performance algorithms for molecular dynamics. *PLoS computational biology*, 13(7):e1005659, 2017.

[7] Jeffrey A Ruffolo, Carlos Guerra, Sai Pooja Mahajan, Jeremias Sulam, and Jeffrey J Gray. Geometric potentials from deep learning improve prediction of CDR H3 loop structures. *Bioinformatics*, 36(Supplement_1):i268–i275, 2020.

[8] Jeffrey A Ruffolo, Jeremias Sulam, and Jeffrey J Gray. Antibody structure prediction using interpretable deep learning. *bioRxiv*, 2021.

[9] Cyrus Chothia, Arthur M Lesk, Anna Tramontano, Michael Levitt, Sandra J Smith-Gill, Gillian Air, Steven Sheriff, Eduardo A Padlan, David Davies, William R Tulip, et al. Conformations of immunoglobulin hypervariable regions. *Nature*, 342(6252):877–883, 1989.
